# Mito-nuclear Incompatibilities and Mitochondrial Replacement Therapy

**DOI:** 10.1101/078329

**Authors:** Adam Eyre-Walker

**Affiliations:** School of Life Sciences, University of Sussex, Brighton, BN1 9QG, United Kingdom

## Abstract

Mitochondrial replacement therapy (MRT) is a human reproductive technology by which the mitochondria of a recipient’s eggs are effectively replaced by those of a donor, potentially eliminating harmful mitochondrial mutations carried by the recipient. However, concerns have been raised that MRT may lead to problems due to incompatibilities between the nuclear genome of the recipient and mitochondrial genome of the donor. Whether this is likely to be a problem is investigated using 226 estimates, taken from the literature, of the effect of replacing the “native” by a “foreign” mitochondrial DNA (mtDNA) from the same species in a variety of animals. In approximately half of the cases (45%), strains with the foreign mtDNA have higher fitness than those with the native mtDNA, and on average the native strains are only 3% fitter. Based on these results it is argued that incompatibilities between the mitochondrial and nuclear genomes are not likely to be a problem for MRT.

---

It has been suggested that incompatibilities between the mitochondrial and nuclear genomes could be a risk factor in mitochondrial replacement therapy (MRT) (Reinhardt *et al.* 2013; Gemmell and wolff 2015; Hamilton 2015; Morrow *et al.* 2015). Incompatibilities between the mitochondrial and nuclear genomes can arise because the vast majority of genes that contribute products to the mitochondrion are located in the nuclear genome (Calvo *et al.* 2016). The risk of these incompatibilities may be higher in MRT because the mitochondrial DNA (mtDNA) is placed into a novel nuclear environment using this process. In contrast, during normal reproduction it is inherited with half of the mother’s genome, a genome the mtDNA has shown itself to be compatible with, because the mother is alive and fertile.

Application of MRT in humans (Craven *et al.* 2010; Paull *et al.* 2013; Hyslop *et al.* 2016; Yamada *et al.* 2016), macaques (Tachibana *et al.* 2009; TAchibana *et al.* 2013) and mice (Sato *et al.* 2005; Wang *et al.* 2014) has generally shown little evidence of deleterious effects which can be traced to mito-nuclear interactions (though see (Hyslop *et al.* 2016)). However, mitonuclear interactions are generally not explicitly tested for, and very few mitonuclear combinations have been investigated. In contrast there is a large body of literature on conplastic strains, in which the mitochondrial DNA from one strain has been placed on the nuclear background of another strain by repeated backcrossing. In some cases the introgression of mtDNA into a strain leads to a loss of fitness; for example, in *Drosophila melanogaster* the introgression of a mtDNA from Brownsville, Texas onto a standard lab background, led to male sterility (Clancy 2008). These deleterious effects have led to the suggestion that there is coadaptation between the mitochondrial and nuclear genomes, and that MRT could be harmful (Reinhardt *et al.* 2013; Gemmell and wolff 2015; Hamilton 2015; Morrow *et al.* 2015). However, there are also cases in which fitness and health are enhanced in conplastic strains; for example the introgression of the mtDNA from the mouse strain NZB/OlaHsd into C57BL/6 led to increase in longevity relative to the C57BL/6 strain with its own mtDNA (Latorre-Pellicer *et al.* 2016).

But what is the average effect? Is there evidence that placing a mtDNA onto a new nuclear background leads to a loss of fitness and health more often than not? To investigate these questions, estimates of the effect of introgressing mtDNA from one nuclear background onto another were compiled from the literature. Only estimates from animals were considered since plant mtDNA has many more genes than animal mtDNA. Studies in which only mitochondrial function had been studied were ignored, since we are ultimately interested in organismal level traits in thinking about human health. Many of the traits that have been studied are probably under stabilising selection in natural populations and hence it can be difficult to judge whether an increase or decrease in the trait is beneficial; we therefore assigned the beneficial direction based on what we might expect if mitochondrial function was seriously compromised; i.e. individuals with very poorly functioning mitochondria might be expected to have a lower chance of survival, develop more slowly, be smaller and have lower fertility. For some traits, we took the direction as that which would be desirable to humans – e.g. increased lifespan and ability to learn – but for some traits, such as exploratory behaviour in mice it was difficult to assign a beneficial direction; these studies were ignored. The dataset comprised 226 estimates, the vast majority of which came from either *Drosophila* species or the beetle *Callosobruchus maculatus* (Supplementary Table S1).

In each case we have the trait value of the line with the nuclear genome and the mtDNA it was isolated with, the “native” strain, and the same nuclear genome with an introgressed mtDNA, the “foreign” or conplastic strain. The magnitude of the effect of introgressing the foreign mtDNA into the strain was quantified as the proportional difference between the lines with the foreign and native mtDNAs (i.e. the difference divided by the mean, calculated such that positive values represent the case when the native line was “better” than the foreign line). Qualitatively similar results were obtained considering the log of the ratio. Only studies using an untransformed scale were included since the proportional difference has little meaning on transformed scales (e.g. the residuals from a regression of trait versus body size).

Unfortunately, the same strains have been measured for many traits and the same trait is often measured at different time-points and temperatures. To reduce the level of non-independence only one time-point and one temperature (the middle value in each case) were included. Nevertheless it should be appreciated that considerable non-independence still exists in the dataset; the same strains have been tested for multiple traits and many of these traits are correlated. As a consequence, no formal statistical analysis was performed on the data. It should also be noted that the vast majority of estimates come from inbred lines; this may exacerbate any interactions between the nuclear and mitochondrial genomes since mutations with large effects tend to be recessive (Simmons and crow 1977). The extent of inbreeding depression in some of these data is evident in the work of Clancy on female longevity in *Drosophila melanogaster* (Clancy 2008). He backcrossed mtDNAs from different localities on to a standard homozygous nuclear background; the mean longevity for mtDNAs from Alstonville, Dahomey and Japan was 27.4 days. However, when those lines were made heterozygous by crossing to another strains of flies, but not the line the mtDNA was sampled from, the mean longevity increased to 38.9 days, a 42% increase.

Taking all the data together there is a slight tendency for conplastic strains to be less fit than native strains (Figure 1); 53% (120) of the comparisons show a negative effect of introgression, 45% (102) show a positive effect, and 2% (4) show no discernible effect (Figure 1). However, the average effect is very close to zero at −2.8% and the majority of effects are small; 57% of the estimates are <10%. If we consider each species individually, for which we have more than 10 estimates, we observe qualitatively similar patterns (Table 1). There is one exception, the copepod *Tigriopus californicus*. In this species the strain with the native mtDNA is always fitter than the strain with the foreign mtDNA, and the average effect sizes is greater than 10%.

**Figure 1.**
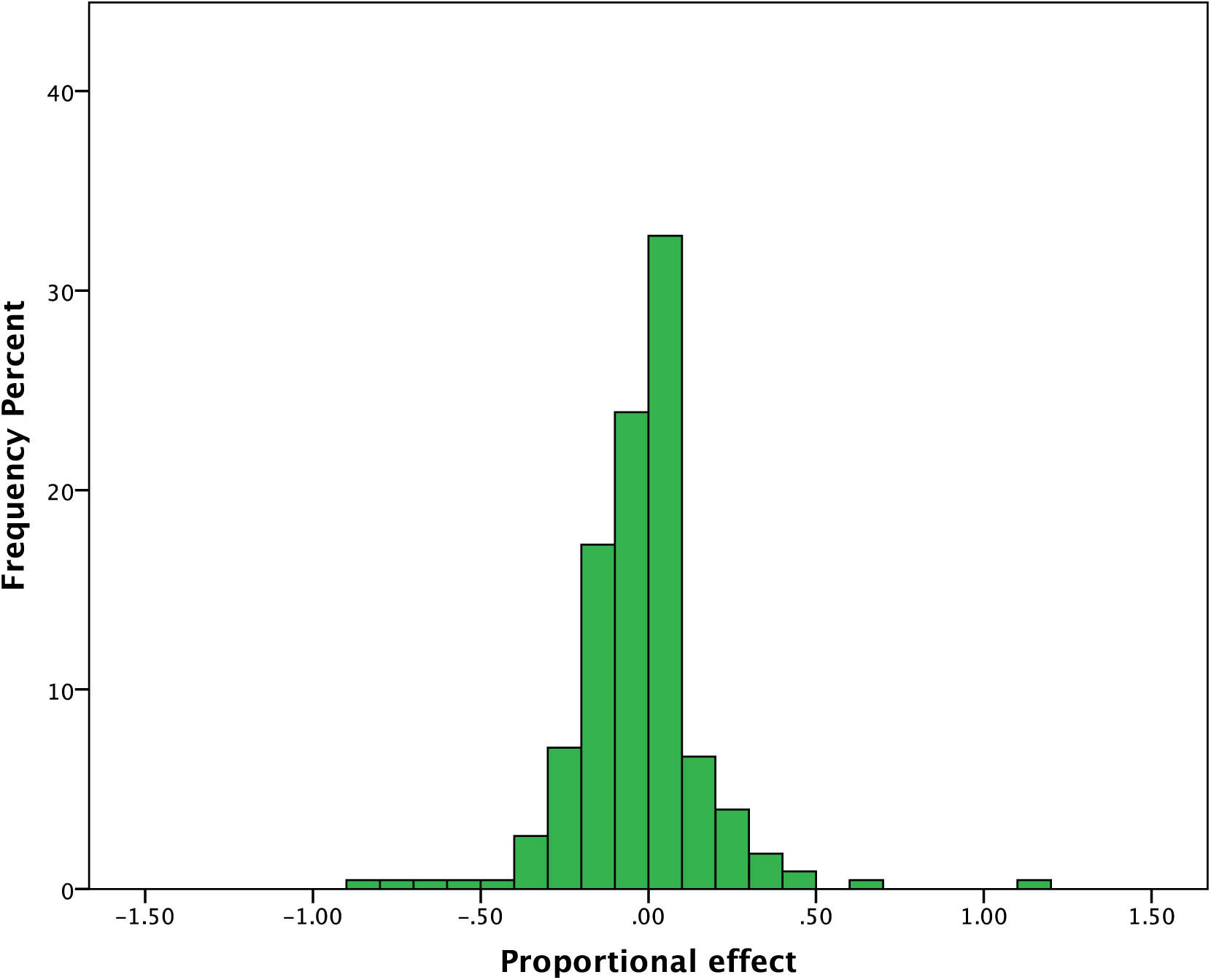
The distribution of proportional effects for 226 estimates taken from the literature. Positive values indicate the strain with the foreign mtDNA is fitter than the strain with the native mtDNA

**Table 1.** The proportion of cases in which the strain with the native mtDNA is better than the strain with the foreign mtDNA, and the average proportional effect (negative values indicate the foreign strain has lower fitness).

| Species | No. of comparisons | Proportion of cases in which Native is better | Average proportional effect (%) |
| --- | --- | --- | --- |
| *Callosobruchus maculatus* | 92 | 0.43 | -1.5 |
| *Drosophila melanogaster* | 66 | 0.60 | -3.4 |
| *Drosophila subobscura* | 32 | 0.44 | -0.1 |
| *Mus musculus domesticus* | 14 | 0.64 | -3.0 |
| *Tigriopus californicus* | 18 | 1.0 | -14 |

It has been hypothesized that the effects of MR might be more apparent in males because mtDNA is not inherited from this sex, and hence mutations that are deleterious in males, but neutral or beneficial in females, can accumulate. There is little evidence that effects of introgression are worse in males; the mean effect in males is −2.7% (79 estimates) and in females it is −2.6% (99 estimates) (Figure 2). There is also little evidence that sampling the mtDNAs from different populations is more deleterious than taking them from the same population – mean effect for between populations is −3.4% (178 estimates) and within it is −0.0% (34 estimates).

**Figure 2.**
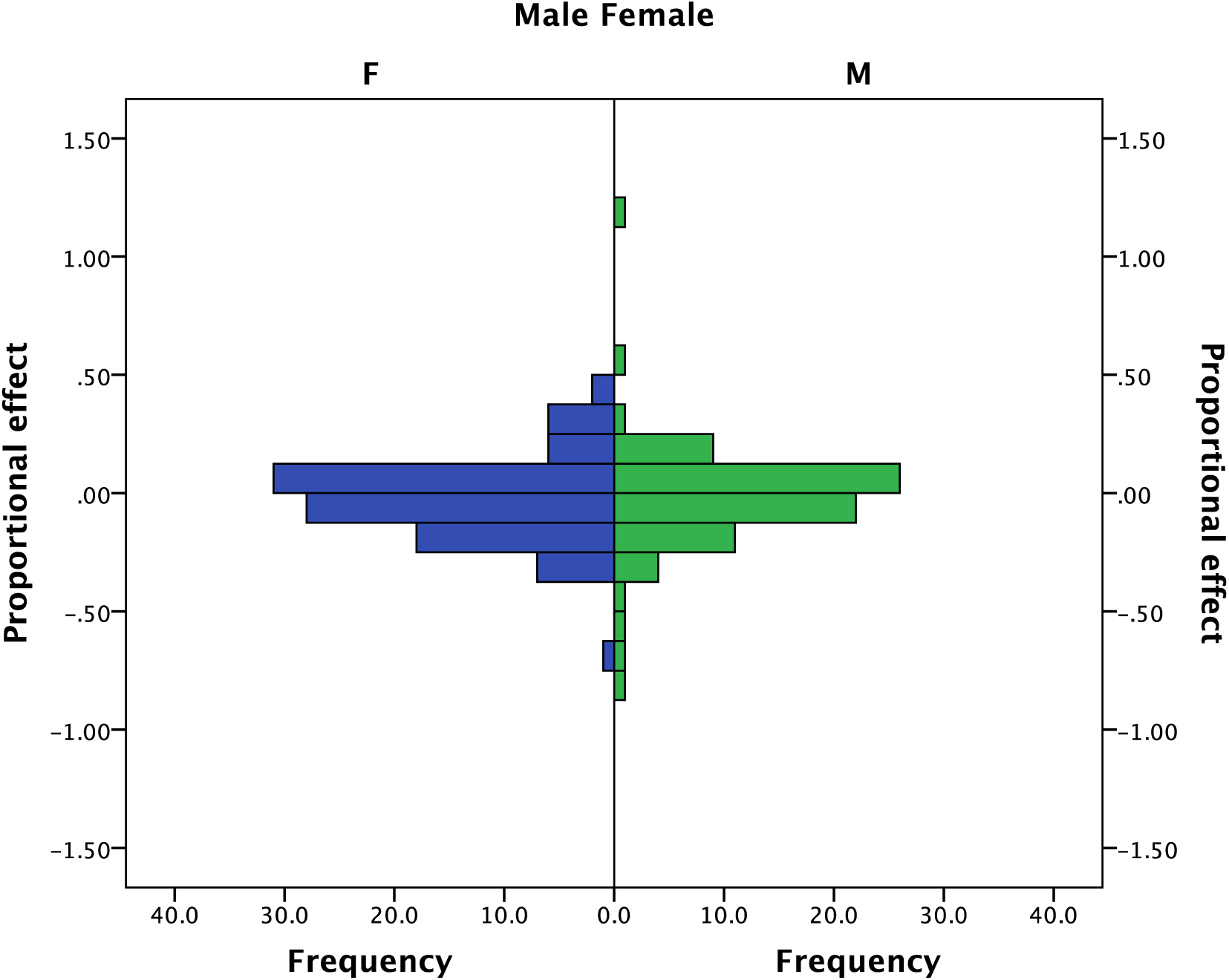
The distribution of proportional effects in males and females.

Overall, there seems to be little evidence that introgressing mtDNA from one strain into another is on average particularly deleterious. This may seem surprising given that the nuclear and mitochondrial genomes are expected to coevolve. However, conplastic strains will only tend to have lower fitness than native non-conplastic strains if there is significant linkage disequilibrium (LD) between the nuclear and mitochondrial genomes. This LD can be established by two mechanisms – selection against deleterious combinations of mitochondrial and nuclear alleles (or selection for beneficial combinations), or divergence between sub-populations. Within a population, selection needs to be strong (>10%) to establish LD, since the rate of recombination is 0.5. Such strong selection is thought to be rare and hence it is unlikely that significant LD between the mitochondrial and nuclear genomes will be established within populations of most species. In contrast, Dobzhansky-Muller incompatibilities can accumulate between populations if there is sufficiently low migration. It is notable in this respect that the species that shows the most consistently negative impact of introgressing a mtDNA from one population to another, *Tigriopus californicus*, shows extremely high levels of population differentiation (Edmands and harrison 2003); FST between the Southern populations of this species, from where the lines were sampled, is 0.81 (Edmands and harrison 2003). In contrast, FST in humans is about 0.05 (Rosenberg *et al.* 2002), similar to the levels observed in *Drosophila melanogaster* (Verspoor and haddrill 2011), which shows no consistent pattern in the effects of introgressing mtDNA into different nuclear backgrounds (Table 1).

In summary, there is little evidence in animals that placing mtDNA onto a new nuclear background, from the same species, is deleterious on average. Hence, there seems to be little reason to expect deleterious mito-nuclear incompatibilities to be more prevalent in individuals born through MRT compared to normal sex, particularly in a species such as humans, which has such low levels of population differentiation.

## Acknowledgements

I am grateful to Damian Dowling for providing his data.

